# Precision functional mapping reveals less inter-individual variability in the child vs. adult human brain

**DOI:** 10.1101/2025.07.21.665760

**Authors:** Damion V. Demeter, Matthew Feigelis, Carolina Badke D’Andrea, Sana A. Ali, Abigail R. Baim, Emily Koithan, Jared Stearns, Salma Zreik, Jonathan Ahern, Sujin Park, Sarah E. Chang, Ryland L. Miller, Jacqueline M. Hampton, Bradley L. Schlaggar, Scott Marek, Evan M. Gordon, Nico UF Dosenbach, Caterina Gratton, Deanna J. Greene

## Abstract

Human brain organization shares a common underlying structure, though recent studies have shown that features of this organization also differ significantly across individual adults. Understanding the developmental pathways that lead to individually unique brains is important for advancing models of cognitive development and neurodevelopmental disorders. Here we use highly personalized precision neuroimaging methods to map brain networks within 12 individual children, ages 8-12 years. We demonstrate fMRI functional connectivity maps that substantially exceed the reliability of traditional techniques, allowing us to measure individual differences after overcoming biases from measurement noise. Children share core functional network topography, with greatest inter-individual variability in association regions, consistent with adult findings. However, children show less between-subject variability than adults, suggesting increasing individual differentiation in brain networks with development. This pediatric precision neuroimaging dataset is publicly available to support future brain development research and provides a high-fidelity foundation for studying individual variation in atypical development.

## Introduction

During childhood and adolescence, brain systems (or ‘networks’), along with cognitive and motor abilities, undergo rapid and significant changes^1–7^. Ontogenetic adaptations that facilitate the transition from early childhood to adulthood, including an increased focus on social behavioral development, accelerated physical growth, and complex changes in the prefrontal cortex, are observed across species^8,9^. The normative developmental trajectory of cognition and motor function has been associated with the selective integration and segregation of various brain regions as part of large-scale functional networks^10–13^. Individual variation around this normative trajectory is likely associated with individual differences in academic achievement^14^, liability to neuropsychiatric illness^15^, and future well-being^16^. Thus, individual differences in functional network organization^17^ may help explain the considerable variability in cognitive function observed among children^7,18^. A deeper understanding of these individual differences may lead to better explanations of variable academic and social outcomes, as well as propensities for neurodevelopmental disorders, and may guide early intervention for at-risk children.

Individual differences in neurodevelopmental trajectories are often studied by testing relationships between brain and phenotypic measures, such as cognitive or clinical features, using group-level neuroimaging approaches (e.g., extracting fMRI signal from group-average defined regions of interest). Large sample datasets, such as the Adolescent Brain Cognitive Development (ABCD) study^19^, provide adequate power to test brain-behavior relationships across thousands of children^20,21^. There has been particular focus on characterizing large-scale functional brain network organization, which is achieved using resting state functional connectivity (RSFC) MRI. These studies have characterized average functional brain networks in children^13,22–24^, providing central tendencies from which to examine generalized changes over development, deviations in psychopathology, and relationships to phenotypic measures.

However, standard amounts of RSFC data per person are noisy at the individual level^25^, hindering the ability to accurately capture reliable individual differences. Moreover, central tendencies inherently blur individual variability in network organization, which can complicate interpretations and impede translational research when used on their own. Group-averages are not necessarily representative of any of the individuals composing them, which has been demonstrated using highly detailed individual-specific representations of functional network structure^26–28^. This limitation is especially problematic in clinical contexts, as individual-specific differences in functional network topography and connectivity could reflect individual clinical phenotypes^29,30^. Specifically, individualized patterns of network organization may be related to differences in disorder presentation and symptom severity that are an important focus for individualized treatment^31^.

Recent fMRI research has seen impressive leaps in the characterization of individual-specific large-scale functional brain networks using a high signal-to-noise ratio precision functional mapping (PFM) approach in adults^28,27,26,32–36^. PFM is the reliable characterization of brain function/organization in an individual person, currently attainable with the collection of hours of non-invasive RSFC data per person over multiple visits^37^. This approach provides a non-invasive method for studies of individual variation in brain function and connectivity in humans and animals alike^38,39^. PFM data provide highly accurate measurements of functional connectivity at the individual level, and have revealed distinct, reliable, and individually unique features in adults despite broadly similar brain network organization^26,30,40^. One feature of the adult brain in particular, “network variants”, defined as locations where an individual’s functional connectivity deviates from the group average, do not coincide with anatomical differences, and are highly stable within an individual across time^30^ and across rest and task states^41^. These findings in the PFM adult literature suggest that individual variation of functional network organization represents stable characteristics of an individual, and therefore, is more likely to provide valuable insights to trait-like features of cognition and behavior.

Given the individual variation in brain organization, a central question arises regarding the developmental pathways that lead to this individual variability. Currently, less is understood about how network variants emerge during development, and the range of individual variation that may be observed during the normative trajectory towards adulthood is unknown. At younger ages, the central tendencies of large-scale organization of functional brain networks generally resembles the organization observed in adults^22,42^, with core network patterns detectable even in neonates^43^. However, there is also evidence of individual variability in network organization in children^24,44^. Moreover, changes in executive function and cognitive control abilities from childhood to adulthood have been associated with changes in network organization^45,46^, supporting the (often implicit) assumption that during development, childhood functional network organization moves toward the “adult-like” brain. A PFM approach comparing child samples to adults can provide deeper insight into the extent to which refinement and specialization in functional organization over development reflects maturation towards a common adult network organization versus deviations from a shared baseline. In addition, given that neurodevelopmental disorders are heterogeneous in their presentation and have been associated with significant differences in brain functional connectivity^47–50^, capturing the neural basis of that heterogeneity and the degree of inter-individual variation in the brain during childhood is especially important for understanding deviations in functional network organization in clinical samples.

The current study introduces a unique, open-access PFM dataset comprising 12 densely-sampled children, each with 1.5 - 6 hours (M=3.3hrs) of unprocessed fMRI data collected over 3 - 12 sessions (M=7.5 sessions), demonstrating feasibility and high within-subject reliability with large quantities of fMRI data per child. We characterize cortical functional network organization in childhood, identifying both shared patterns and individually-unique features across individual children. Furthermore, we quantify the extent to which the topography and connectivity of functional networks varies across individuals and compare these metrics to an established adult PFM dataset^26^. Finally, we compare inter-individual variability across age-groups, revealing that children exhibit less inter-individual variability in brain functional connectivity compared to adults. PFM datasets not only provide appropriate data for discoveries of inter-individual variation of brain function, but also have the potential to help establish gold standards for reliability and allow for tests of validity with results observed in samples containing standard amounts of fMRI data collection.

## Results

### Collection of PFM data is feasible in a child sample

The collection of PFM data in a pediatric sample can be challenging^51^ due to children’s lower tolerance for repeated or prolonged scan visits and exhibiting greater head motion than adults^52^. Here, we demonstrate the feasibility of obtaining high-quality (i.e., low motion) PFM data from 12 children and introduce the child Precision Functional Mapping (cPFM) dataset. These children ranged in age from 8.2 - 11.9 years (M=9.9, SD=0.9), included six males and six females (gender identity was not collected), and included three children with a reported neurodevelopmental disorder diagnosis and one child with an incidental finding of a benign cyst in the medial frontal cortex. One additional child was recruited and screened but withdrew from the study prior to scanning due to claustrophobia. Each participant completed 3 - 12 MRI visits, during which T1w and T2w structural MRI scans and up to three 10-min resting state fMRI scans were collected per visit. After strict motion censoring to mitigate artifactual effects of head motion^53–56^ (See Methods), each participant retained between 1 and 5.5 hours of usable resting state fMRI data (M=3 hours), with 64 - 95% of total data retained per participant (M = 77%, Figure 1A). With these high quality data, participants demonstrated considerable within-participant stability, indicated by the high correlation of visit-to-visit RSFC within-participant and low visit-to-visit RSFC between-participants (Figure 1B). Thus, the cPFM dataset demonstrates feasibility of obtaining a minimum of 1 hour, and up to 5.5 hours, of low-motion, usable resting state fMRI data across at least 3 scan sessions in children. Table 1 reports additional scan and demographic information.

**Figure 1.**
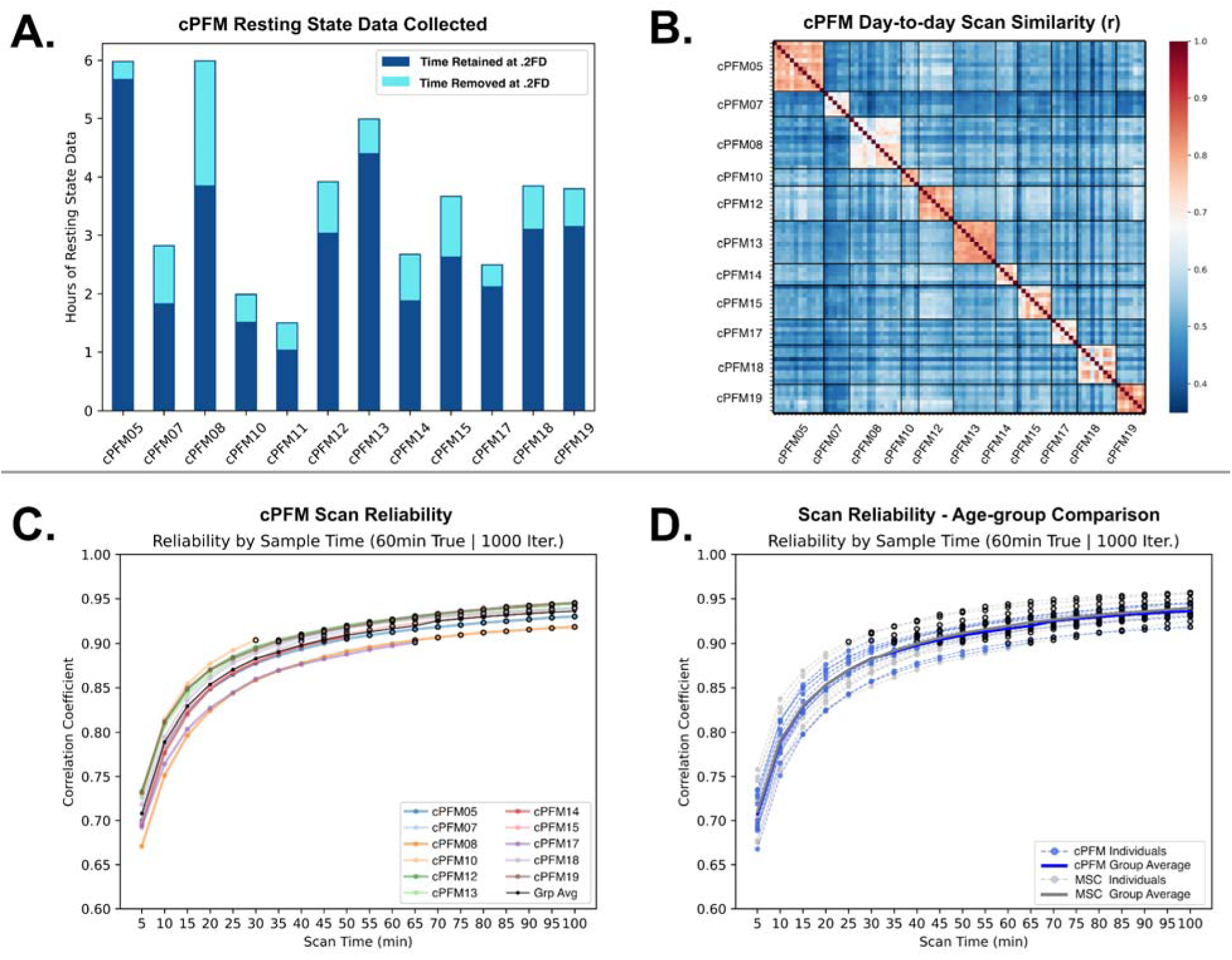
Resting state fMRI data summary, similarity, and reliability for child PFM data. **A.** Hours of resting state data collected for each individual, with colors indicating how much time was retained versus removed due to motion (framewise displacement (FD) threshold = 0.2mm). **B.** Visit-to-visit similarity within and between participants, such that each participant’s RSFC data from one visit was correlated to their other visits’ data and all other participants’ data from each visit. **C.** Reliability curves of RSFC data for each participant in the cPFM dataset (each participant shown in a different color) with at least 90 minutes of scan time; circles are bolded when reliability ≥ 0.9. **D.** Reliability curves for each participant in the cPFM dataset (blue) and in the MSC adult dataset (gray). Group average of reliability curves for cPFM and MSC are displayed with solid lines. Abbreviations: cPFM = child Precision Functional Mapping; Grp Avg = Group Average, MSC = Midnight Scan Club. Note: cPFM11 is included in A only, due to less than 90 minutes of remaining data after motion censoring

**Table 1.** Participant Demographic Information.

| Participant Demographic Information |  |  |  |  |  |  |  |  |  |
| --- | --- | --- | --- | --- | --- | --- | --- | --- | --- |
| Child Precision Functional Mapping Dataset |  |  |  |  |  |  |  |  |  |
| ID | Sessions | Total Time Collected | Time ≤ 0.2 FD | Age at First Scan | Sex | IQ | Race | Ethnicity | Diagnosis |
| cPFM05 | 12 | 5:59:42 | 5:40:14 | 9.2 | M | 122 | White | NH | None |
| cPFM07 | 6 | 2:49:52 | 1:49:42 | 10.62 | M | 97 | White | NH | None |
| cPFM08 | 12 | 5:59:42 | 3:51:12 | 9.88 | M | 118 | White | NH | None |
| cPFM10 | 4 | 1:59:54 | 1:31:12 | 10.12 | M | 122 | Black, White | NH | None |
| cPFM11 | 3 | 1:29:56 | 1:01:36 | 9.17 | M | 114 | Black, White | NH | None |
| cPFM12 | 8 | 3:55:15 | 3:02:07 | 9.8 | F | 134 | White | NH | Graves Disease |
| cPFM13 | 10 | 4:59:45 | 4:24:19 | 9.66 | F | 126 | Asian, White | NH | Benign brain cyst |
| cPFM14 | 5 | 2:41:38 | 1:52:37 | 8.24 | M | 110 | White | NH | TS, ADHD |
| cPFM15 | 8 | 3:39:49 | 2:37:51 | 11.9 | F | 102 | White | NH | TS |
| cPFM17 | 6 | 2:29:53 | 2:07:12 | 10.13 | F | 137 | White | NH | None |
| cPFM18 | 9 | 3:51:23 | 3:05:43 | 10.58 | F | 114 | White | NH | TS |
| cPFM19 | 7 | 3:48:06 | 3:09:18 | 9.44 | F | 109 | White | NH | None |

NH = Non-Hispanic, TS = Tourette Syndrome

### RSFC is highly reliable with child PFM data

To quantify the amount of data required for reliable estimation of cortical functional connectivity, we conducted an iterative split-data reliability analysis^26,28^ (See Methods). Figure 1C shows that reliability reached a correlation of 0.9 for the majority of participants when using 45-50 minutes of “test” data, measured against an independent held out 60 minutes of “true” data. Additionally, cPFM reliability curves show minimal but non-negligible increases in reliability beyond 90 minutes of data. The group average of the reliability curves in the cPFM dataset also follows a trajectory similar to the group average reliability curve for adult PFM data (Midnight Scan Club; MSC). Reliability curves compared across children and adults did not show group differences and the group average reliability curves were nearly fully overlapping for the cPFM and MSC datasets (Figure 1D).

### Individualized functional networks show broad similarities as well as individual features across children

Functional network topography for each child was identified using data-driven community detection^26,57^ (See Methods). Each participant exhibited individual-specific network organization comprising 12 canonical functional networks: visual (VIS), somatomotor hand (SM Hand), somatomotor face (SM Face), somatomotor foot (SM Foot), auditory (AUD), default mode (DMN), cingulo-opercular (recently termed the action-mode network^58^ and referred to as CON/AMN hereafter), frontoparietal (FPN), dorsal attention (DAN), language (LANG), salience (SAL), and contextual association (CA) networks (Figure 2A; All views shown in Supplementary Figure S2). Children shared broadly similar topographic organization of their functional networks, with between-participant Normalized Mutual Information (NMI) of network assignment = 0.44. At the same time, individual differences were apparent, with variability across individuals in specific features of each network. The reliability of these network maps was evaluated through a split-half procedure as in^42^, finding an average split-half reliability of network assignment of 0.66 NMI between halves within the same individual (Figure 2C).This result of higher within-subject NMI than previously reported is likely due to the increased reliability of densely-sampled PFM data. Split-half reliability values revealed a bimodal distribution corresponding to participants with total post-processed and motion-censored scan time either above or below 2 hours (i.e., more data corresponded to larger NMI), supporting the benefits of larger quantities of per-person data.

**Figure 2.**
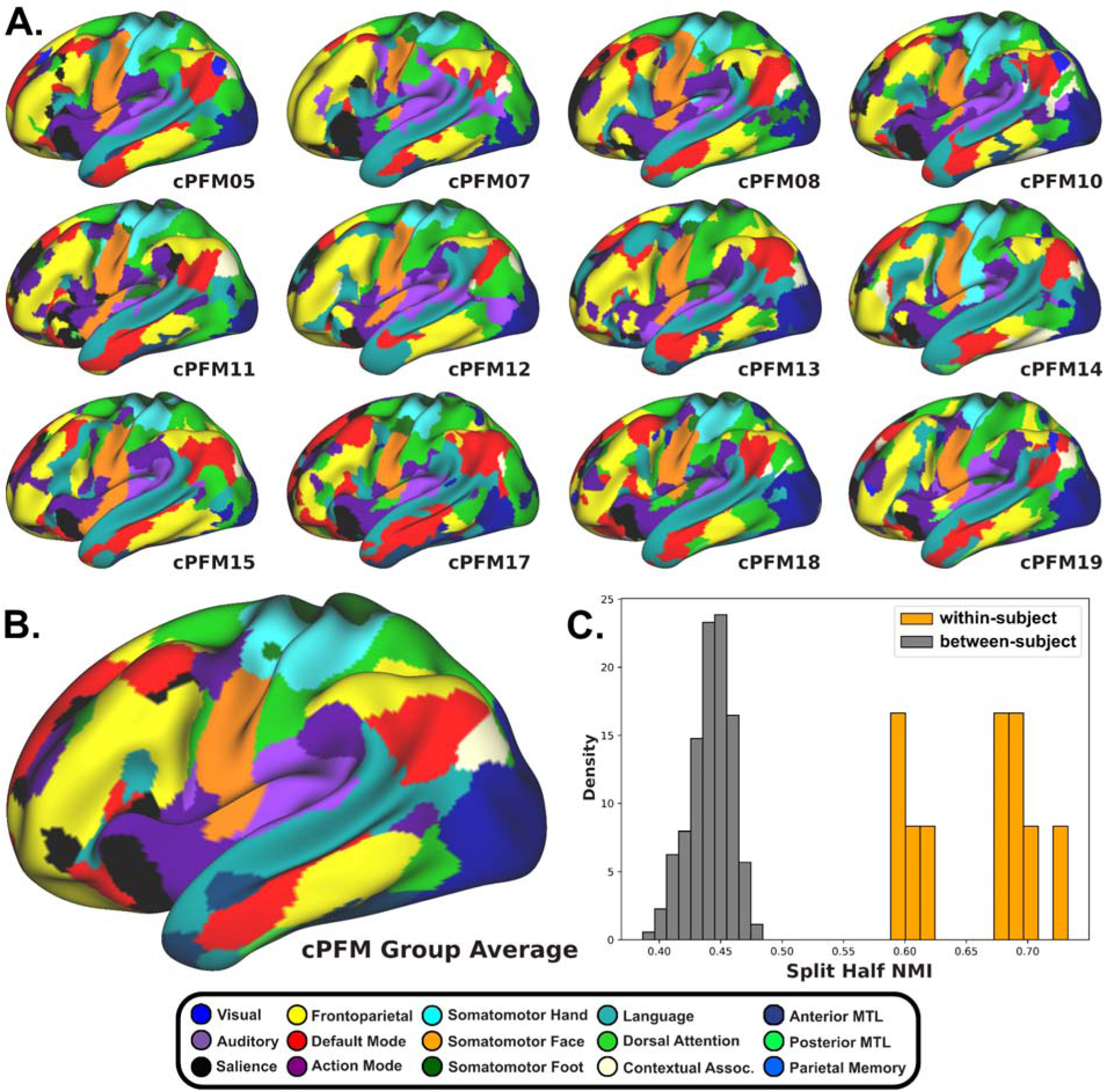
Individual child cortical functional networks. **A.** Individual cPFM participant cortical functional network topography displayed on the left hemisphere, lateral surface (all views shown in Figure S2). **B.** cPFM group average network topography. **C.** Split half NMI comparing within-subject and between-subject network assignments. Bimodal distribution of participants for within-subject estimates coincides with amount of data (more data = higher NMI).

To evaluate the consistency of functional network assignments among children, we generated density maps for each network, which show the proportion of occurrences of each given network at every cortical vertex across participants. As depicted in Figure 3, large consistency was evident for each network with the largest regions of consensus occurring in somatomotor and medial vision regions. Consistent with results from adult data^26,40^ and probabilistic network maps from child ABCD data^42^, we observed canonical attributes in all children, including DMN features in bilateral angular gyri, FPN in the lateral prefrontal cortex, and a dorsal-ventral distinction between somatomotor foot, hand, and face regions. Further, core regions of each network showed the largest consistency among the group, with regions on the edge of each network displaying the largest variability.

**Figure 3.**
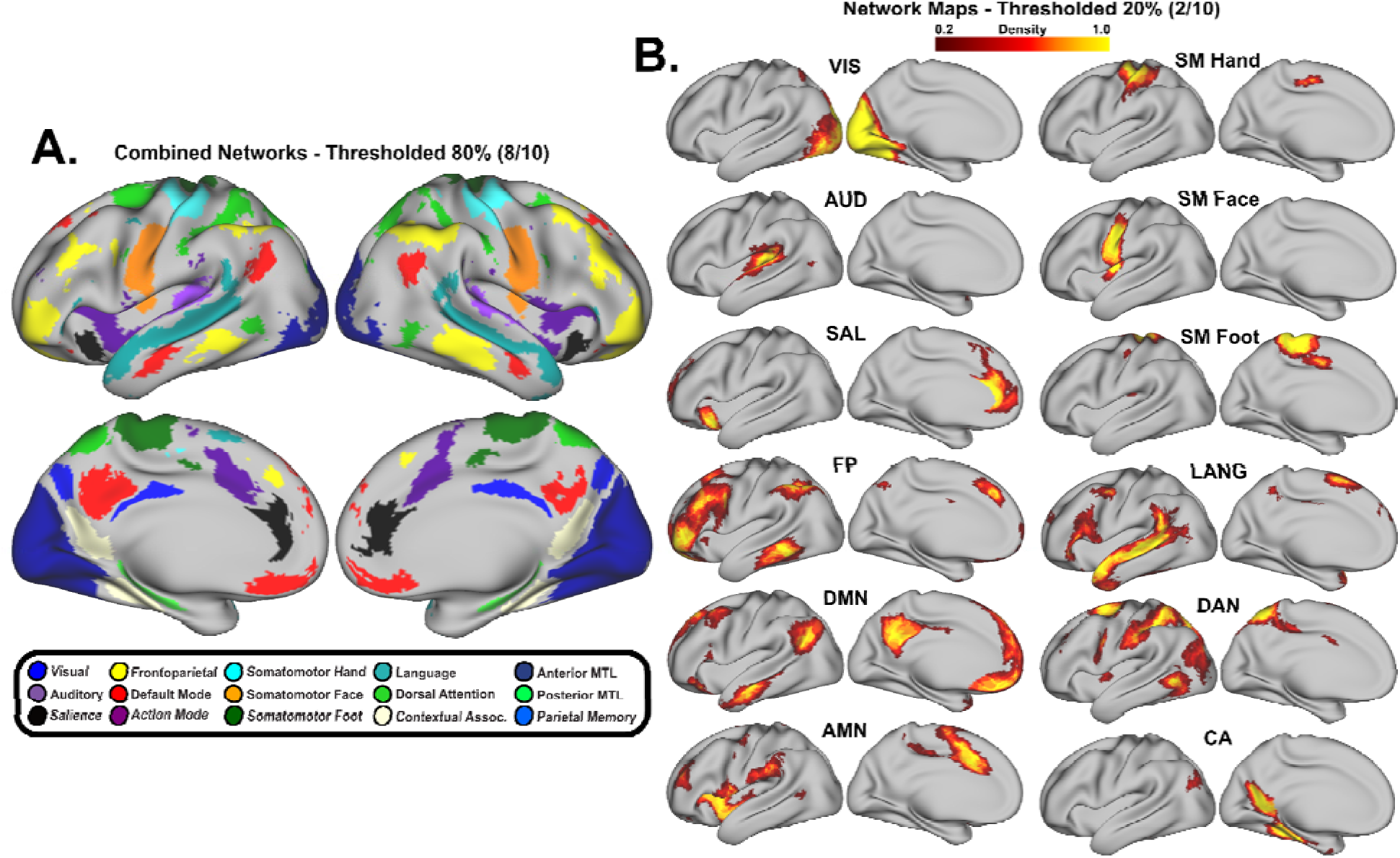
Regions of high network similarity across individual children. Vertex-wise density maps. **A.** Colored vertices indicate where 80% of individuals have the same network assignment. **B.** Density maps for each of 12 canonical functional networks. All network maps are shown with an applied threshold of 20% (vertices where that network was assigned for at least 2 of 10 cPFM participants). Group figures exclude cPFM11 due to < 90 minutes of low-motion data and cPFM13 due to a benign brain cyst in medial frontal cortex.

### Individual children show network variants - distinct brain regions with individually variable functional connectivity

To investigate individual variability in RSFC, cortical network variants – regions of the brain with functional connectivity that significantly differs from a group average – were identified for each child. Following similar methods previously applied to adult PFM data^30,41^, individual RSFC was calculated for all cortical vertices in each child and compared to a group average derived from 185 children from the ABCD study (selected based on data collection at the same site as the cPFM dataset, and for having sufficient amounts of low-motion RSFC data; demographics in Table S1). All cPFM children exhibited cortical RSFC variants, with variant locations and expected network associations differing across individuals. RSFC variant locations in the children were generally qualitatively similar to those reported in adults^30,41^, located primarily in the bilateral frontal cortex, temporo-occipito-parietal junction, and along the cingulate gyrus (regions of group overlap are displayed in Figure 4A). Variants in the cPFM group were more common in the right hemisphere than the left hemisphere, showing similar lateralization to that previously reported in adult PFM data^59^. Split-half similarity analyses of functional variants within- and between-individuals (Figure 4C) showed high similarity within individuals (quantified by the Dice coefficient; average split-half Dice = 0.744) and significantly lower (p < 0.001) similarity between individuals (average between-individuals Dice = 0.291). The cPFM group also showed the highest agreement of variant location across participants in cortical regions assigned to the canonical FPN and LANG functional networks (Figure 4B,D).

**Figure 4.**
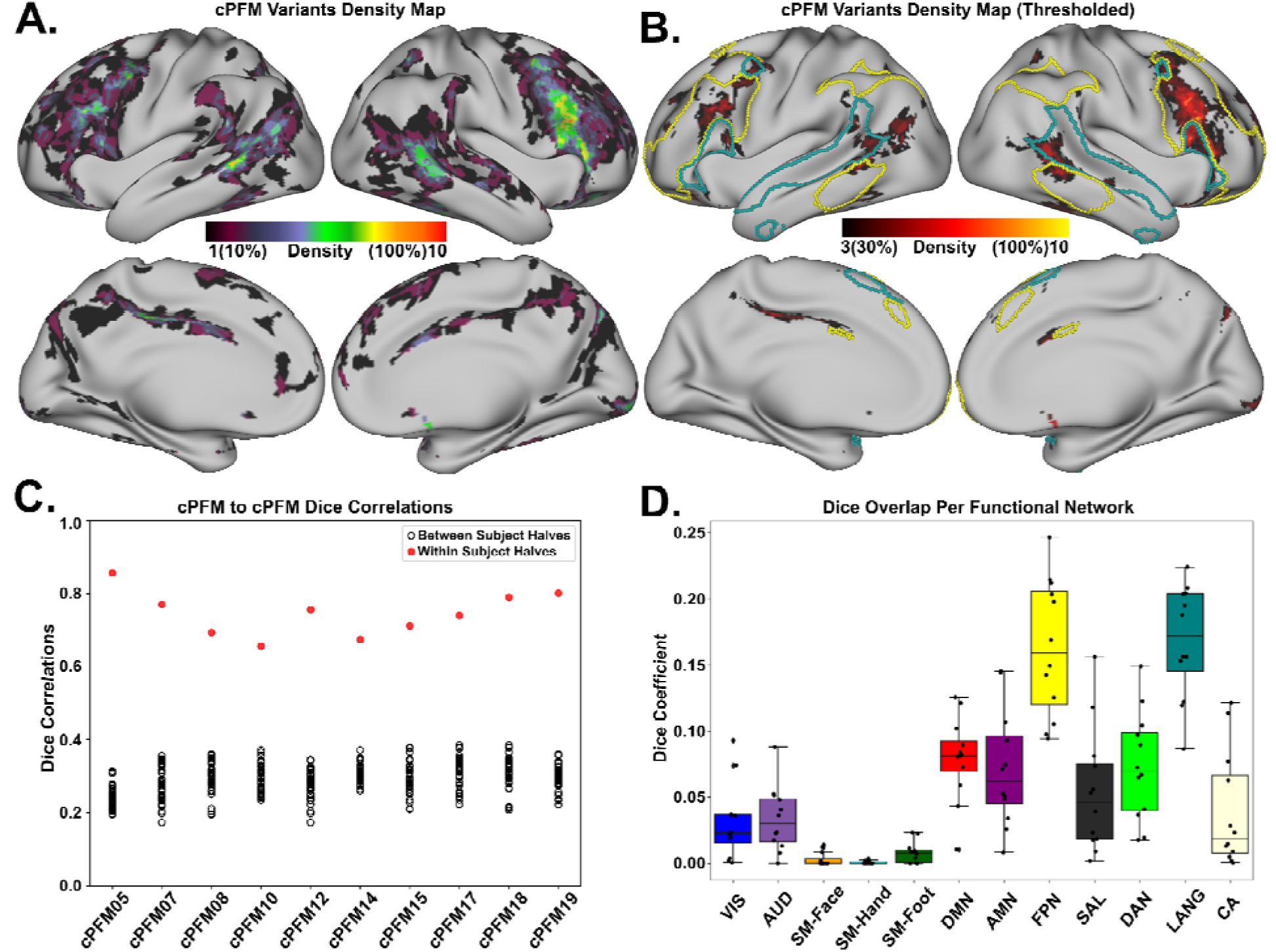
Network variants: regions of variation in individual children. **A.** cPFM group variant density map displays the percent of individual children with variants in a given cortical location. **B.** cPFM group variant density map thresholded to a minimum of network variants observed in 3 participants overlaid onto outlines of functional networks that overlapped most with variant locations (FPN in yellow, LANG in teal). **C.** Split-half Dice coefficient correlations within (red circles) and between (black circles) all cPFM participants. **D.** Boxplots depicting the variants’ overlap with each canonical RSFC functional network across the cPFM group. (Group figures exclude cPFM11 due to < 90 minutes of low-motion data and cPFM13 due to a benign brain cyst).

### Inter-individual variability is lower in children compared to adults

Having established an understanding of the similarity and variability in individual-specific functional network organization among children, we next compared the children (cPFM) to a set of adults with PFM data (MSC). Inter-individual similarity of functional network organization was quantified at the whole-cortex vertex-wise level and at the regional parcel-wise level for these comparisons.

We quantified the similarity between individuals in the same age cohort (e.g., child-to-child) by computing the correlation of whole-cortex vertex-wise RSFC data between all pairs of participants within each age group (Figure 5A). As expected from the previously described similarity in functional network organization, inter-individual similarity was fairly high in the children (mean z(r) = 0.61; sd = 0.05). In the adults, inter-individual similarity was also fairly high (mean z(r) = 0.52; sd = 0.03). Yet, the inter-individual within-group similarly in the adults was significantly lower than that in the children (p < 0.001, Figure 5B), suggesting inter-individual variability in network organization increases with age.

**Figure 5.**
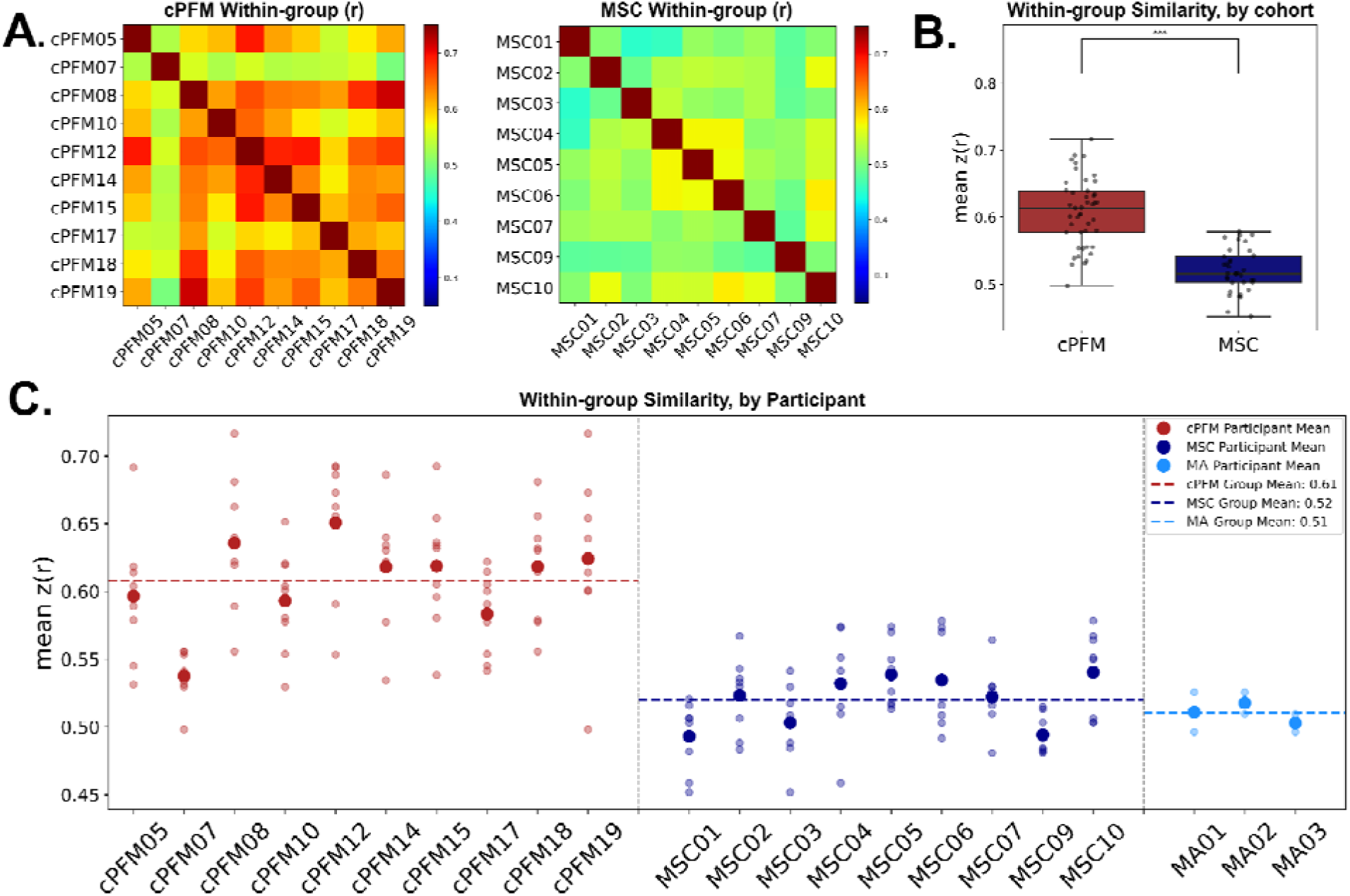
Inter-individual similarity in functional connectivity is higher among children than among adults. **A.** cPFM and MSC participant-to-participant similarity matrices comparing whole-cortex functional connectivity within each age group. **B.** Mean within-group similarity by age cohort shows significantly higher similarity in the children compared to the adults (p < 0.001) **C.** Within-group similarity values for each individual. On the right, scanner type and sequence matched adults (MA01, MA02, MA03) had identical scanning parameters to the cPFM group.

To control for total scan time, as the MSC individuals have more data per person than the cPFM individuals, we conducted a control analysis sampling an identical amount of scan time and frame count from each participant, and found similar results (parcel-wise results shown in Supplementary Figure S3). Despite the adults having more scan time than the cPFM group on average, the children had higher inter-individual similarity than the adults. To control for the potential influence of MRI scanner and sequence differences, we ran the same analysis on three adult participants from a separate study^60^ (referred to here as “MA” for matched adults) that used the same scanner and sequences as the cPFM participants (Figure 5C). Notably, two of these adults were also part of the MSC dataset, allowing for a small within-person, cross scanner comparison (MSC02-MA01 mean z(r) = 0.6; MSC06-MA02 mean z(r) = 0.75). The within-group similarity by participant (Figure 5C) strongly suggests that scanner type or sequence differences did not drive the observed age-related differences between cPFM and MSC datasets.

To investigate which cortical regions displayed the highest inter-individual similarity, we computed within-group similarity maps at the vertex level for each age group. In both cohorts, within-group similarity was largest in primary somatomotor and medial visual regions, the insula, precuneus, and posterior cingulate cortex (Figure 6A,B). When calculating the difference between these maps (child - adult), the child cohort had overall larger within-group similarity values than the adult cohort across the somatomotor, superior parietal, temporal, and medial prefrontal cortex. (Figure 6C; between-group similarity depicted in Supplementary Figure S5).

**Figure 6.**
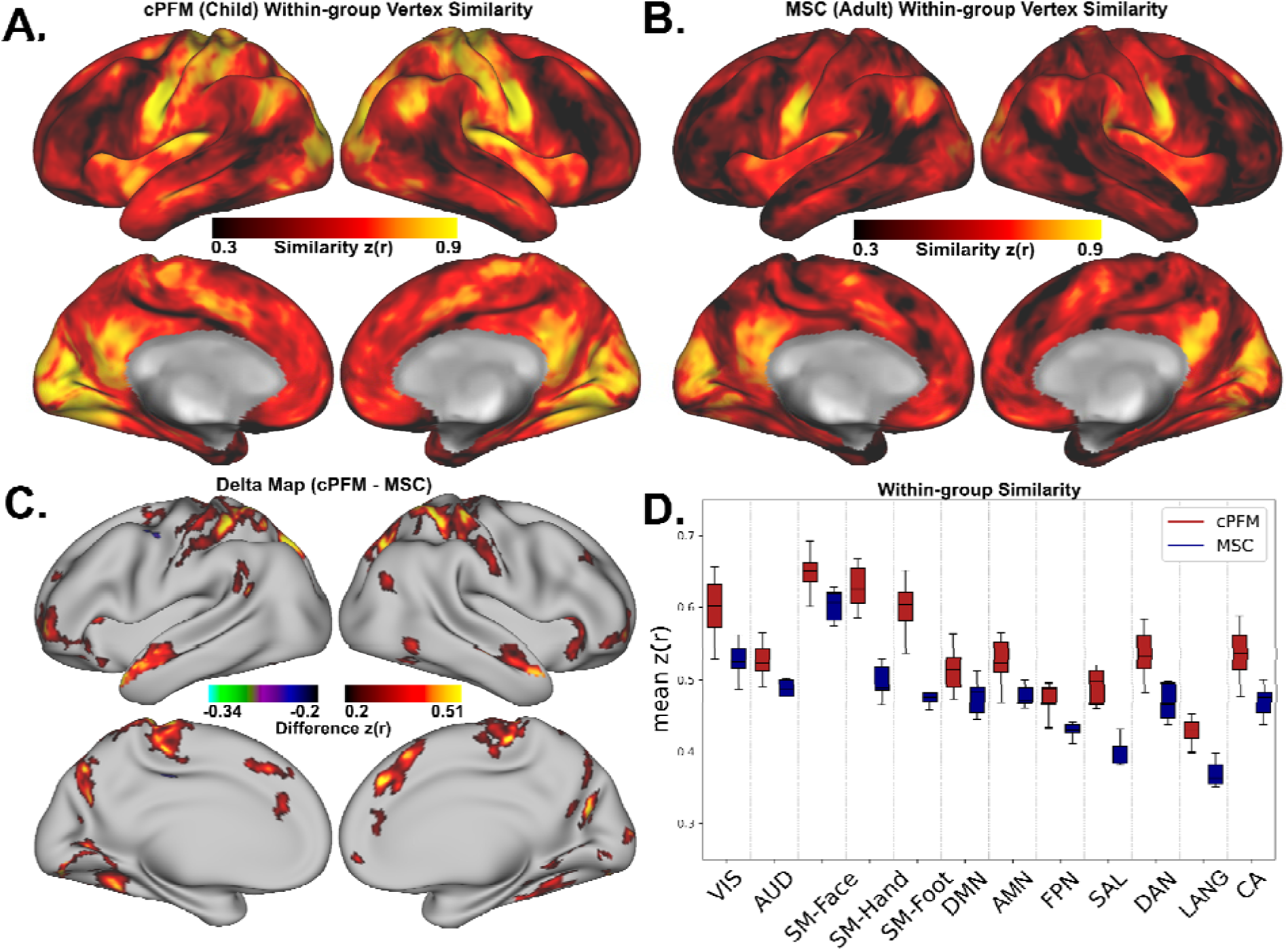
Within age-group similarity across the cortex. Vertex-wise within-group similarity map for **(A)** the child group, and **(B)** the adult group. **C.** Difference map displays where within-group similarity is higher for the children than for the adults (50mm^2^ cluster size threshold applied for figure). **D.** Within-group similarity broken down by individual-specific network identity labels.

We further grouped vertex values by individual-specific functional network identity at each vertex and found that child within-group similarity values were significantly larger in every functional network after permutation testing (FDR p < 0.05, Figure 6D). These results demonstrate higher within-group similarity of the child cohort across the whole cortex and not selectively confined to specific networks.

## Discussion

Here, we present and release a unique fMRI dataset of densely sampled children, demonstrating feasibility and high reliability of functional network measures at the individual- child level. Using these highly precise individual maps, we show that while exhibiting some individual differences, children are more similar to each other, i.e., less individually variable, in their functional brain network organization than adults. These results suggest individualized refinement towards unique brain organization from childhood to adulthood, rather than movement from more individually variable organization in childhood to a stable “adult-like” mature state.

Given the evidence showing that the core patterns of functional network organization are already in-place in neonates^43^, there is likely a large genetic component expressed in utero that establishes human baseline brain organization^61,62^. Unique coactivations of brain regions and Hebbian-like processes may then selectively strengthen or weaken functional connectivity^63^, potentially leading to increased inter-individual variability during the lifetime. Cortical regions whose borders are broadly genetically predetermined and comprise continuous gradients in the embryonic cortex, are later refined through extrinsic sensory input contributing to the development of functional areas and large-scale networks postnatally^64,65^. Our findings suggest that this refinement leads to individualization, in which individuals deviate more and more from the central tendency with maturation. Hence, we propose a “starburst” model of neurodevelopment in which the intrinsic functional organization of the brain begins more similar and then differentiates with development to refine each person’s unique brain. This refinement occurs in every functional network, suggesting maturation is a global process impacting the whole brain rather than being limited to a specific network or region. Indeed, age is associated with brain network integration and segregation across various sensory and control systems^66^, with RSFC-age relationships seemingly distributed across the whole brain^67^. Further, functional connectivity of several different functional networks exhibit relationships with measures of cognition and executive function^17,24,66^. Therefore, neurodevelopmental changes throughout childhood and adolescence are not limited to any specific system, but rather, occur throughout the cortex.

One may have predicted more inter-individual variability in children given the variation in cognitive abilities and developmental stages in the age range of children sampled here, 8- to 12- years-old (M=9.9, SD=0.9)^7^. As individual differences in cognitive abilities have a direct association with trajectories of brain development and growth^66,68^, it would follow that these varying developmental stages would coincide with greater inter-individual variability in brain network connectivity. Indeed, there is evidence that brain activation patterns during emotion processing are more inter-individually similar in older than younger children^69^. However, our findings of more inter-individual similarity in children compared to adults suggest that the inter-individual variability in intrinsic functional brain organization, which has been shown to reflect stable features of an individual^29^, may develop differently from task-evoked activations.

Alternatively, these findings may highlight inter-individual variation that is better captured with densely-sampled, high-reliability data. Either way, our results indicate that as we age, the intrinsic organization of functional brain networks becomes more variable across our same-age peers. Indeed, inter-individual variability in brain function has been shown across species and scales^66,69^, including evidence for activity-dependent individual differentiation in neurophysiology^70^. Hence, increased inter-individual variability in functional network organization with development may be due to the accumulation of unique experiences, cognitive strategies, behaviors, or environmental and genetic factors prior to adulthood^71^.

It is also possible that other biological factors contribute to the development of individual differences in network organization. For example, specific age-related changes in vasculature^72,73^, myelination^74^, and cortical thinning^75^ may play a role in the level of individual variation in functional connectivity that is observed in adults at different ages. Further, the effects we observe with functional connectivity data may, in fact, relate to downstream effects of inter-individual variation of gyri and sulci at full maturity reported in large mammals^76^. Future longitudinal studies across multiple age ranges will provide a more complete understanding of the factors leading to increased inter-individual variation in brain networks as we age.

The current study also replicates and extends multiple findings on adult individual-specific functional brain organization to a pediatric population. First, as was reported in adults^26^, we demonstrated that high-fidelity, functional network mapping of brain networks in a pediatric sample can reveal individual-specific organizational variability of brain networks. We observed individually-specific features of functional brain network organization in childhood that were obscured in previous studies using common group-average approaches. While group-average analyses are important for understanding population level brain organization, individual features of functional networks are necessarily obscured^26,77^, and the ability to measure inter-individual variation is reduced^78–80^. Second, we demonstrated that probabilistic mapping of functional networks created from PFM data reveals a stable and generalizable central tendency and high consistency of RSFC network topology as shown in adult PFM work^40^, as well as large sample, standard collections of childhood fMRI data^42^, such as the Adolescent Brain Cognitive Development (ABCD)^19^ study. We delineated functional brain networks for each child that consisted of 12 canonical networks, each with common regions of consensus among the cores of these networks. This finding supports previous work showing a stable group effect of functional brain networks^29^ and the well-accepted idea that “adult-like” brain organization is already in place in childhood, and may even be present at birth^43^. Third, sensorimotor networks were less individually variable than association (e.g., cognitive, attention) networks, an effect that can be measured with standard amounts of fMRI data per person^81,44^. Yet, the stable cores of these networks that allow for some estimation of inter-individual variability may not sufficiently reflect the nuance and precision of person-specific variation of these networks that requires PFM datasets. Lastly, we found that cortical network “variants” are reliably identified during childhood using PFM data, as previously demonstrated in adult PFM work^30^. Regions of inter-individual variability, specifically where an individual’s functional connectivity differed substantially from the group (i.e., variants), were located primarily in association cortex as well as along the borders of each network. These network variants were stable within an individual, unique to that individual, and located in largely similar regions as previously reported in adults^30,41^. Variants with the most agreement across cPFM participants were found in regions assigned to the canonical FPN and LANG functional networks.

It is possible that methodological or group factors may contribute to differences between children and adults, posing some limitations to the current study. The primary datasets used here (cPFM and MSC) had different fMRI acquisition settings, which led us to include three adults from a different PFM study (“MA” for matched adults) with identical acquisition settings to the cPFM dataset. These adults still demonstrated greater inter-individual variability amongst their same-age peers than that of the children. In addition, the adult data were collected on a structured schedule with scan sessions occurring at approximately the same time of day, whereas the child data were collected at different times of day per each family’s availability.

Despite the impact of introducing potential variability due to time of day effects, we still observed greater within-group similarity of functional connectivity within the children relative to the adults. Another potential difference between the groups could be neuropsychiatric diagnoses or other medical conditions. Three of the children in the cPFM dataset had at least one neuropsychiatric disorder diagnosis, one had an autoimmune disorder, and one had a benign medial frontal cyst (incidental finding). Again, we would expect this variability in the child sample to increase inter-individual variability in brain function, which is inconsistent with the decreased inter-individual variability we found in the children. Still, future PFM collections focusing on neuropsychiatric disorders are needed to better understand how different phenotypes affect inter-individual variability. Recent work has demonstrated that PFM data can identify inter-individual differences in brain connectivity related to depression^82^, and future work using child PFM data may help identify phenotypes for childhood neuropsychiatric disorders. Additionally, the adult datasets consisted of more total usable data than the child dataset. For several analyses, we matched the amount of data between the groups, which did not change the results. Moreover, we would reasonably expect that less data, which leads to reduced within-subject reliability, would cause greater inter-individual variability. Yet again our results are inconsistent with this expectation, suggesting that scan time did not confound the findings. Finally, the child-to-adult comparison was necessarily cross-sectional. Therefore, we note that the effects reported here demonstrate *differences* between children and adults, which does not necessarily indicate *changes* with development. Future longitudinal PFM studies can illuminate developmental changes directly.

The current study demonstrates the feasibility of collecting large quantities of high-quality resting state fMRI data per person in a pediatric sample, and we are publicly releasing the child PFM (cPFM) dataset as a resource for other researchers to use in future analyses. Although pediatric populations are generally considered more logistically challenging than adults for fMRI data collection due to concerns of in-scanner head motion and lower tolerance for repeated scanning^52^, we successfully collected an average of 3 hours of low-motion resting state fMRI data per child from 12 children, age 8-12 years, over 3 to 12 scanning sessions each. Despite an expected higher level of in-scanner motion for this age group, we retained an average of 75% of each participant’s data after stringent motion correction. Moreover, we achieved reliable estimates of functional connectivity of a quantity and quality comparable to previous adult PFM datasets^26^. Specifically, high within-subject reliability (test-retest r > 0.9) of vertex-wise cortical functional connectivity was reached with approximately 45-50 minutes of high-quality, low motion data. It should also be noted that the cPFM dataset includes 3 participants with neurodevelopmental disorders. While this collection is not sufficient in numbers to effectively compare network organization between typically developing children and those with neurodevelopmental disorders, the general topography and organization of functional networks were similar across child participants.

Together, this study provides evidence for the feasibility of dense sampling and a PFM approach in a pediatric sample, demonstrates inter-individual similarity and variability of functional brain networks during childhood (and compared to adults), and suggests a trajectory of brain network refinement that begins similar in childhood and becomes more individually unique as we age, consistent with a “starburst” model of neurodevelopment. The results presented here and the release of the cPFM dataset for future work provide a foundation for understanding how brain networks undergo such refinement and may inform how inter-individual differences in brain networks relate to phenotypic differences. It is also important to note that studies using PFM datasets should work in tandem with large N studies^83,84^, such as the ABCD study^19^, to provide insight to both findings of high generalizability, as well as individual variability that is obscured through group-average approaches.

## Acknowledgments

We would like to acknowledge the incredible work done by all research staff involved in the collection and curation of this dataset (past and present) and the families that dedicated their time to participate in this study.

## Author Contributions

Conceptualization: D.V.D, M.F, D.J.G; data curation: D.V.D., M.F., C.B.D., S.A., R.L.M, J.S; data processing: D.V.D., M.F.; formal analysis and visualization: D.V.D., M.F.; methodology: D.V.D., M.F., C.G., N.U.F.D., E.M.G., D.J.G.; software: D.V.D., M.F., E.M.G.; writing – original draft, D.V.D., M.F., and D.J.G; writing - review and editing: D.V.D., M.F., S.A., A.B., E.K., S.Z., J.A., S.P., S.C., R.M., B.J.S., S.M., E.M.G, C.G., and D.J.G; supervision: D.J.G.

## METHODS

### Participant demographics

Thirteen children were recruited to participate in this study; one withdrew prior to scanning due to claustrophobia. Hence, the Child Precision Functional Mapping (cPFM) dataset reported here comprises 12 children, ages 8-12 years old (6 female, 6 male). Three children met diagnostic criteria for neurodevelopmental disorders, and a benign brain cyst was discovered as an incidental finding in one child. Detailed demographic and diagnostic information is reported in Table 1. Children were recruited from the Washington University community and from databases of previous participants who were willing to be contacted again for future studies.

Parents or legal guardians provided informed consent, and all child participants provided assent. The Washington University School of Medicine Human Studies Committee and Institutional Review Board approved this study.

The Midnight Scan Club (MSC) dataset of ten healthy young adults (5 female, 5 male; mean age = 29.1±3.3 years, range 24-34) was used as an adult comparison PFM dataset. Detailed demographic information is detailed in Gordon et al. (2017). Three participants (two of whom were also in the MSC dataset; ages 25, 27, 35 years) from a different study were used to test for scanner and MRI sequence effects. Demographic information for those adults is detailed in Newbold et al. (2020).

### Neuroimaging acquisition

All cPFM participants were scanned at Washington University School of Medicine on a Siemens Prisma 3T MRI scanner with a 64-channel head coil. Foam padding was applied around the head for participant comfort and to mitigate head motion during scans. Verbal feedback was given between scans to ensure participant comfort and to provide feedback on participant motion. Real-time motion analytic software – Framewise Integrated Real-time MRI Monitoring (FIRMM)^52^ – was used to track participant motion during scans.

Participant data were collected over 3 to 12 visits (M=7.5). At least one T1-weighted structural MPRAGE sequence (TR=2500ms, TE=2.9ms, FOV=256x256, voxel resolution=1x1x1mm) and one T2-weighted structural image with turbo spin echo sequence (TR=3200ms, TE=564ms, FOV=256x256, voxel resolution=1x1x1mm) were collected across visits and used for preprocessing. Up to five 10-minute echo-planar sequence functional resting state scans (TR=1100ms, TE=33ms, flip angle=84°, MB factor=4, 54 axial slices, voxel resolution=2.6x2.6x2.6mm) were collected per visit. During resting state scans, participants were instructed to view a white fixation cross on a black background, stay awake, and lie as still as possible.

The adult Midnight Scan Club^26^ dataset was downloaded from www.openneuro.org (doi:10.18112/openneuro.ds000224.v1.0.4) in unprocessed NIfTI (Neuroimaging Informatics Technology Initiative) format. fMRI data specifics can be obtained from previously published material^85^.

The adult data used to test for scanner and MRI sequence influences (MA01-03) can be downloaded from www.openneuro.org (doi:10.18112/openneuro.ds002766.v3.0.2).

Preprocessed fMRI data used for the analyses included in this study were obtained from authors of the original study and full fMRI preprocessing specifics (which closely matched those listed below) can be found in the original published work^60^.

### Imaging data preprocessing

All MRI data were preprocessed in-house, using a public release of the DCAN-Labs abcd-hcp-pipeline^86^. Additional FMRIB Software Library^87^, Freesurfer^88^, and Connectome Workbench^89^ commands and custom MATLAB^90^ scripts were also used for preprocessing and data analysis. The DCAN-Labs abcd-hcp-pipeline follows the primary steps of the Human connectome minimal preprocessing pipeline^91^, followed by additional resting state focused preprocessing steps, informed by best practices in the field.^92–97^.

The resting state functional connectivity focused preprocessing steps included: (1) de-meaning and de-trending of data; (2) general linear model “denoising” of signal related to white matter, cerebral spinal fluid, whole brain (global) signal, and six directions of motion plus their derivatives; (3) temporal band-pass filtering (0.008Hz < f < 0.09Hz); (4) respiratory motion filtering^98^ (5) and motion censoring which excluded frames exceeding a framewise displacement (FD) of 0.2mm. Additionally, retained frames were required to be in clusters of at least 5 contiguous, below-FD-threshold frames. Registration steps and denoising are each done in a single pass to mitigate the reintroduction of noise^97^.

All resting state data were then mapped to an MNI-transformed midthickness 32k fs_LR surface mesh^99^ and spatial smoothing was applied via geodesic Gaussian smoothing (6mm FWHM, 2.55 sigma) to create the final CIFTI dense timeseries file used for analyses.

### Reliability curves

Within-subject RSFC reliability curves were calculated to visualize the reliability of scan data across multiple visits and identify the quantity of RSFC data needed for within-participant reliability to gain little value from additional data collection. Reliability curves were calculated for all cPFM participants with greater than 1 hour and 30 minutes of data (n=11) using the following method. For each individual, resting state timeseries were extracted using 333 previously defined cortical parcels^100^. First, we created a baseline or “true” RSFC correlation matrix using 1 hour of data, pseudo-randomly sampled across all scan visits of that individual. This ensured that data from all scan visits were represented in each correlation matrix. A “test” correlation matrix was then created by pseudo-randomly sampling the remaining RSFC data, again, distributed across all scan visits, in 5-minute increments and the “true” and “test” matrices were correlated. The data used to create the “test” correlation matrices was increased until all remaining data was used. This process was repeated 1000 times and the average, per-participant, reliability plots for the cPFM dataset are shown in Figure 1C.

### Vertex-wise individual-specific network identification

Individual-specific functional network organizations were identified using the Infomap community detection method^101^, similar to the methodologies presented in Power et al.^57^ and Gordon et al.^26^. In short, we computed pairwise Pearson r correlations among the BOLD time series across all cortical vertices, generating a correlation matrix of dimensions 59,412 x 59,412.

Subsequently, this matrix was thresholded across a range of densities spanning from 0.1% to 5%. For each threshold, data-driven community assignments were found using the Infomap algorithm. To attribute putative network identities to each community at each threshold, we utilized a template matching procedure. The Jaccard index was used to calculate the spatial overlap of each community with a set of independent networks (Supplementary Figure S1) derived from a cohort of 7,316 children in the Adolescent Brain Cognitive Development (ABCD)^19^ study (See supplementary Table S1 for ABCD 7,316 group demographics). An assignment was given to the best matching network if the Jaccard index was at least 0.1 (less than 0.1 overlap was not given an assignment to prevent poorly fitting matches). To consolidate assignments across sparsity thresholds, a consensus network assignment was derived by retaining the network identity of a vertex at the sparsest threshold where it was successfully assigned to a known group network.

### Probabilistic maps

We conducted a probabilistic/density map analysis similar to the approach of Dworetsky et al.^40^. For each cortical vertex, we tallied the incidences of each network assignment across all children. cPFM11 and cPFM13 were removed from this and subsequent analyses due to having < 90 minutes of low-motion fMRI data and a benign mPFC cyst; Supplement Figure S4), respectively. Following this step, every network map underwent normalization by dividing it by the total number of individuals within the corresponding age group. A combined network map was created by finding the most common network occurrence at each vertex across the children.

### Functional Network Variant Identification

#### Group average RSFC comparison data

Cortical network variants (regions that show strong dissimilarity to the group average functional connectivity) were identified using methods previously applied to adult datasets^30,41^. First, each individual’s connectivity matrix was compared to an age-appropriate RSFC group average. For the cPFM participants, the comparison group average connectivity matrix was created from 185 (83 F) children from the Adolescent Brain Cognitive Development (ABCD)^19^ collection, ages 9-10.8 years old (M=10.2yrs), and did not include both individuals from any sibling or twin pairs (See supplementary Table S1 for RSFC comparison group demographics). All participants included in this group average were required to have at least 5 minutes of resting state data post motion censoring at 0.2mm FD (M=13m 55s). Additionally, all retained frames were required to be within groups of at least 5 contiguous, below-FD-threshold frames. In order to best mitigate methodological differences, the 185 ABCD participants were scanned at the same location (Washington University School of Medicine) as the cPFM set, and data were preprocessed with the same DCAN-Labs abcd-hcp-pipeline^86^ and smoothed with a matching smoothing kernel of 6mm FWHM.

For the adult comparison (MSC) participants, the WashU 120^102^ group average was used, matching methods in the previous adult work using MSC data. fMRI preprocessing, motion censoring, and spatial smoothing were the same as described above.

### Cortical network variant calculation

Cortical network variants for each participant were identified by correlating the functional connectivity of each surface vertex of a given individual with the functional connectivity of the matching vertex in the group average data from the corresponding age group. This procedure resulted in an individual-to-group average spatial correlation map for all 59,412 cortical surface vertices. Cortical regions with the lowest 10% of correlations that also consisted of at least 30 contiguous surface vertices were then binarized and labeled as a functional network variant. The binarized variant maps for all participants were then used for all subsequent variant analyses.

### Calculation of inter-individual variant map similarity

Spatial overlap of participant variant maps was quantified using the Dice similarity coefficient^103^. The Dice similarity coefficient is a spatial overlap index, ranging from 0 which indicates no spatial overlap of two binary sets, to 1 indicating complete overlap of two binary sets. Binary variant maps for all participants created in the previous step were used to calculate inter-individual variant map dice coefficients across both age groups to quantify the age-related similarity of functional connectivity variants of children and adults.

### Functional network similarity

We calculated pairwise correlations between whole-cortex resting state functional connectivity at the vertex level, for each pair of participants in each age cohort. An RSFC matrix was constructed for each participant using 59,412 cortical surface vertices. Subsequently, we determined the correlation between the upper triangular components of the RSFC matrix for each participant and those of all other participants.

### Within and between age-group similarity maps

To investigate which brain regions exhibit higher similarity in children compared to adults, we created an average spatial correlation map for each age group. The functional connectivity of each cortical vertex of each participant was correlated with the connectivity at that vertex with every other individual in the age group. This procedure resulted in an individual-to-individual spatial correlation map for all 59,412 cortical vertices for each pair of individuals in each age group. These maps were then averaged in each group to create within-age-group similarity maps. This analysis enabled us to identify cortical regions with large similarity within their respective age groups and how within-group similarity differed between the child and adult cohorts. To further investigate the latter, a difference map was created by taking child within-group similarity minus adult within-group similarity maps.

An identical approach was taken to create the between-age-group similarity map, except individual-to-individual spatial correlation maps were created from each child to each adult pair, and then the average of these maps was used to create the between-group similarity map.

The vertices were further categorized into 12 canonical functional networks: visual (VIS), somatomotor hand (SM Hand), somatomotor face (SM Face), somatomotor foot (SM Foot), auditory (AUD), default mode (DMN), cingulo-opercular action-mode (CON/AMN), frontoparietal (FPN), dorsal attention (DAN), language (LANG), salience (SAL), and contextual association networks (CA).

## Data Availability

Data from the Midnight Scan Club (MSC) dataset are publicly available at https://openneuro.org/datasets/ds000224.

Data used in the preparation of this article were obtained from the Adolescent Brain Cognitive Development (ABCD) Study (https://abcdstudy.org). ABCD consortium investigators designed and implemented the study and/or provided data but did not participate in the analysis or writing of this report. This manuscript reflects the views of the authors and may not reflect the opinions or views of the NIH or ABCD consortium investigators. The ABCD data repository grows and changes over time. The ABCD data used in the analyses of this report can be found under the NDA Study DOI: 10.15154/1522676. DOIs can be found at https://nda.nih.gov/general-query.html.

Correspondence and requests for additional materials should be addressed to Deanna J. Greene

