## Supplemental Figures for "Precision functional mapping reveals less inter-individual variability in the child vs. adult human brain"

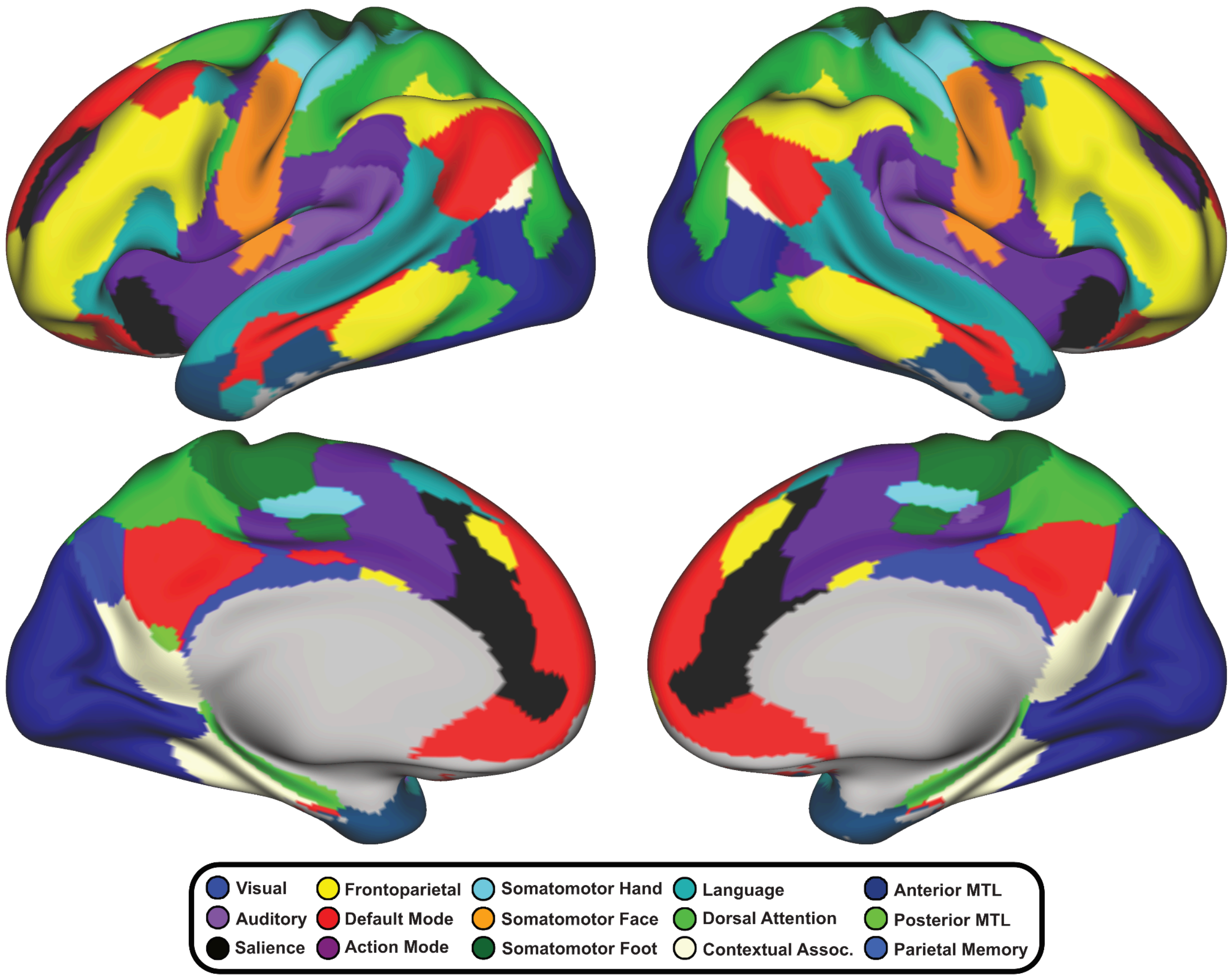

***Supplementary Figure S1 -*** ABCD 7,316 Template
The group cortical network organization template used in child analyses within this study was created from 7,316 9-10.9 year-olds from the publicly available Adolescent Brain Cognitive Development (ABCD) study.

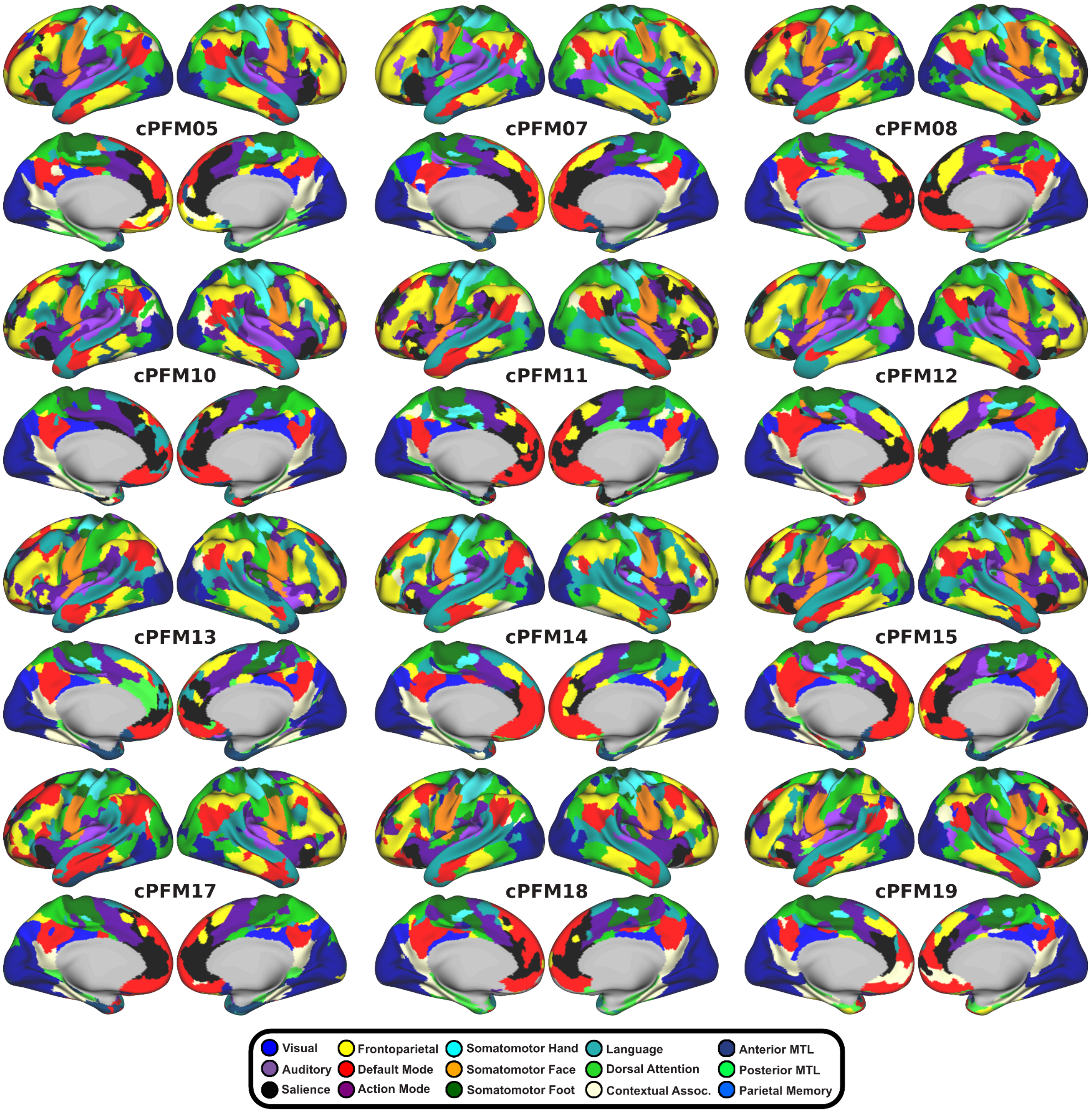

**Supplementary Figure S2 -** All views of cortical functional networks for the cPFM dataset (related to Figure 2).

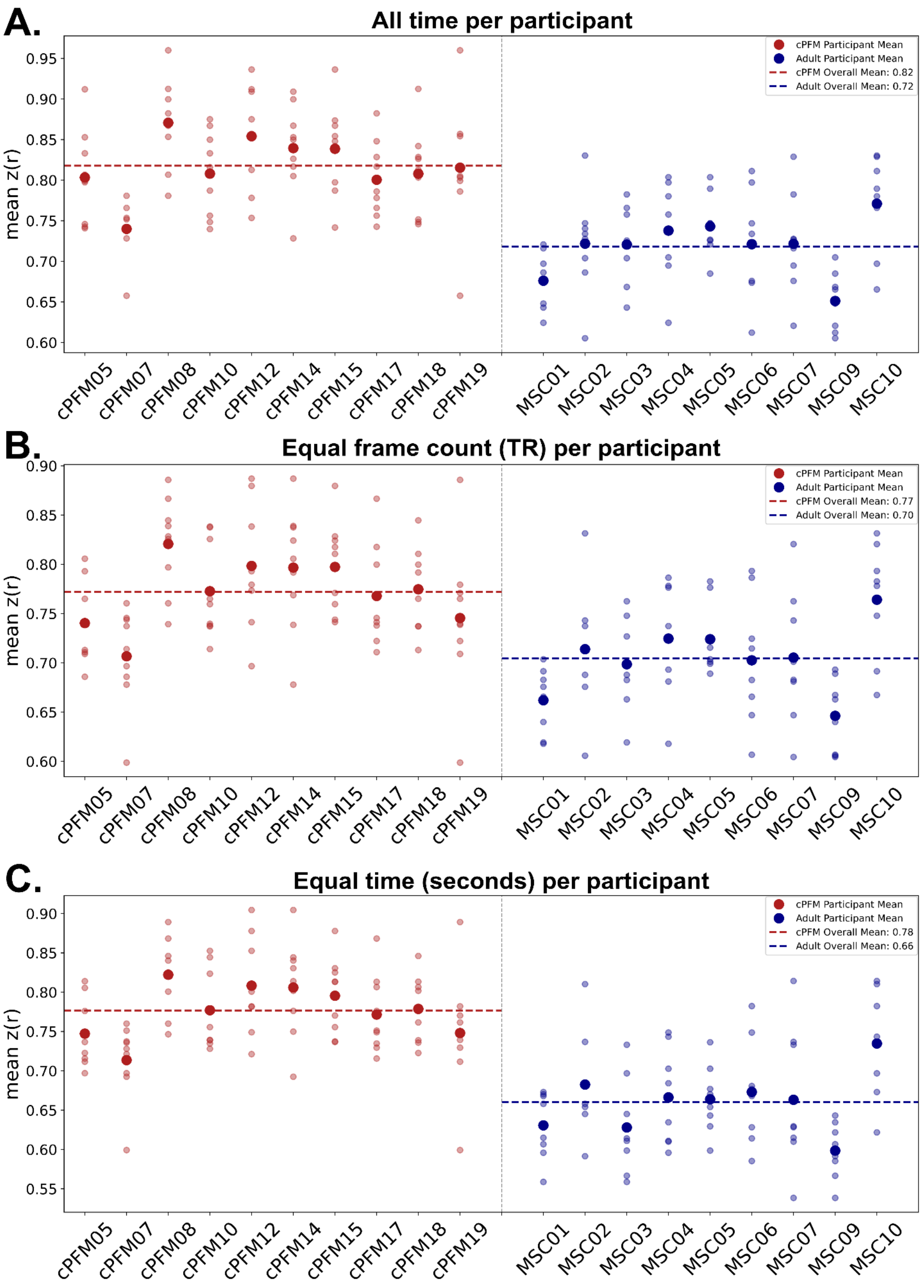

**Supplementary Figure S3 - Within-age group inter-individual similarity accounting for amount of per-person data**

As cPFM and MSC groups were collected using different TR values (1.1TR, 2.2TR respectively) and had different scan durations for each participant, we reran the inter-individual variability analysis using equal frames (i.e., TRs) and time (in seconds) per participant. **A.** Parcel-wise within-group similarity values for each participant including all data retained after preprocessing. **B.** Within-group similarity values using the same number of frames per participant. We reduced each individual's data to the minimum frame count across both groups (frame count = 4,572). **C.** Within-group similarity values using the same amount of scan time per participant. We reduced each individual's data to the minimum scan time across both groups (minutes = 91.2). (Figure excludes cPFM11 due to < 90 min low-motion data, and cPFM13 due to a benign brain cyst).

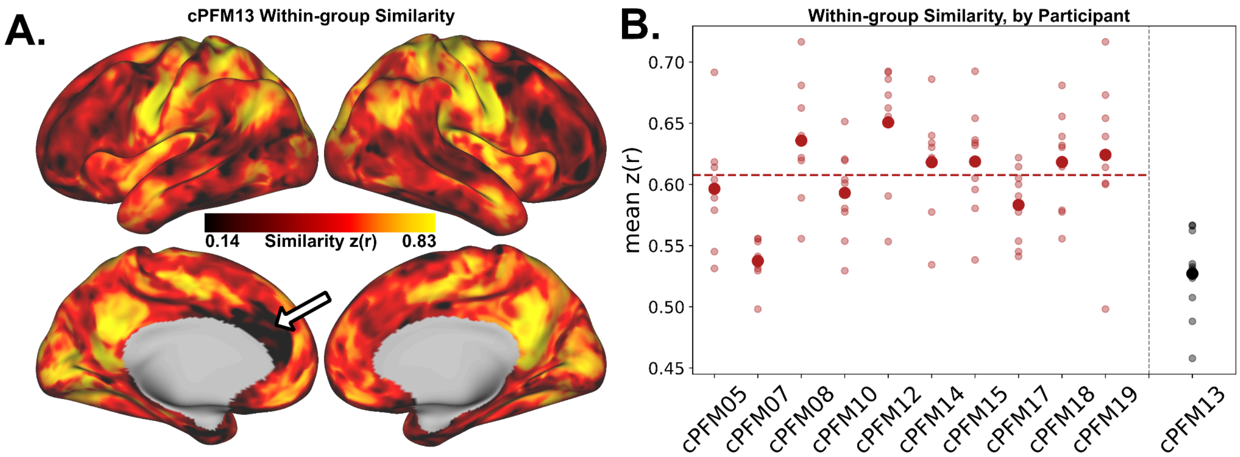

**Supplementary Figure S4 - Lower inter-individual variability in child with benign mPFC cyst**

Related to Figure 5: Within-group similarity values, including participant with benign cyst in medial prefrontal cortex (cPFM13). Arrow depicts the cyst location.

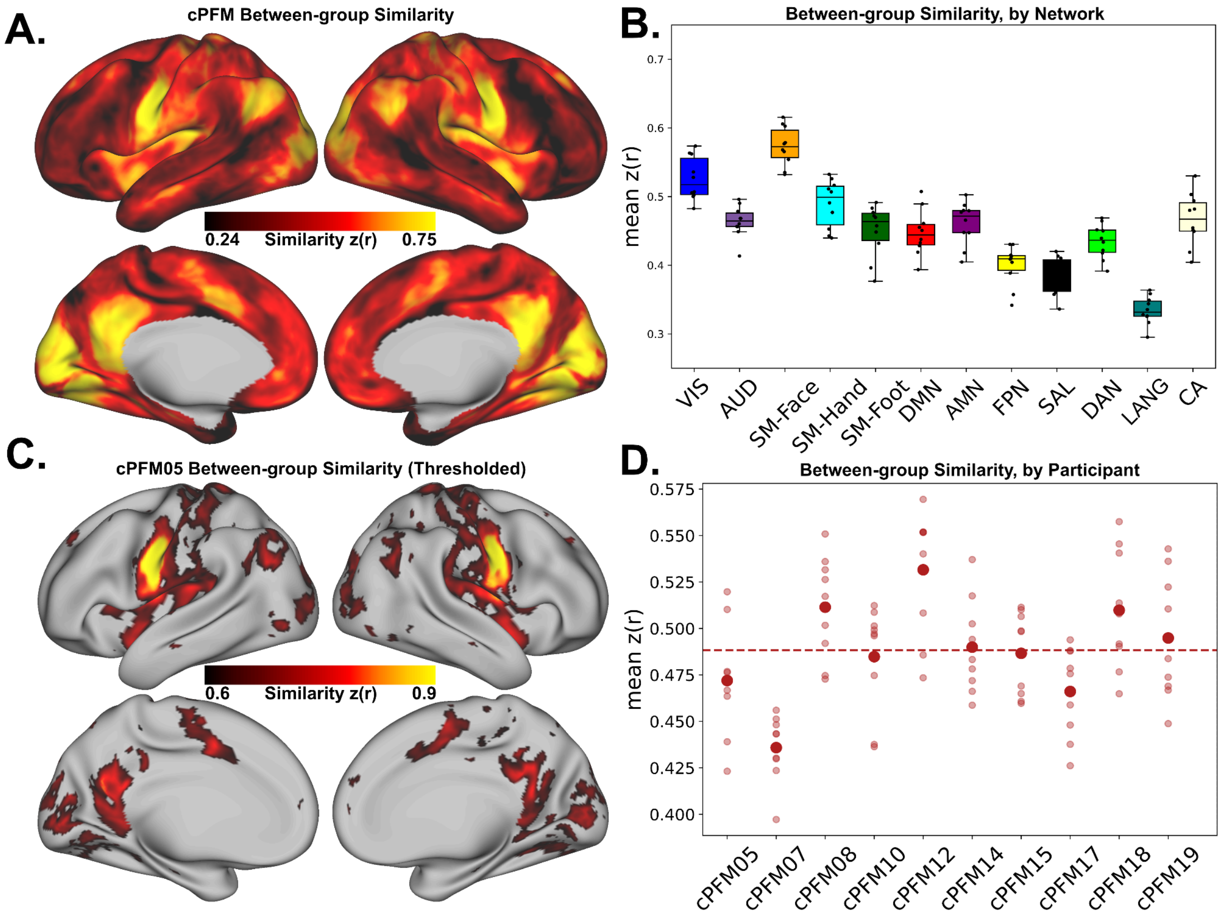

**Supplementary Figure S5 - Between age-group similarity map demonstrating regions of age invariance**

**A.** Between age-group similarity map created from averaging across all child-to-adult comparisons. **B.** Between-group similarity broken down by network label. **C.** Individual-specific between-group similarity map for a representative participant cPFM05. **D.** Between-group similarity values for each child compared to each adult in the MSC dataset.

**Table S1 - RSFC Comparison Groups Demographics**

| Group RSFC Comparison Data Demographics | |
|---|---|
| **ABCD 185 group** | |
| Participants | 102 Male (55%), 83 Female (45%) |
| Age Range (years) | 9-10.9 years |
| Age (Mean) | 10.2 years |
| Mean Scan Data | 13 min 55 sec |
| Handedness | 139 R (75%), 16 L (4%), 38 Ambi (21%) |
| **Race and Ethnicity** | |
| White | 74 (40%) |
| Black | 84 (45%) |
| Hispanic | 5 (3%) |
| Asian | 0 |
| Other | 22 (12%) |
| Not Reported | 0 |
| **ABCD 7,316 group** | |
| Participants | 3,667 Male (50.1%), 3,649 Female (49.9%) |
| Age Range (years) | 9-10.9 years |
| Age (Mean) | 9.9 years |
| Mean Scan Data | 15 min 13 sec |
| Handedness | 5,891 R (81%), 501 L (7%), 917 Ambi (12%), 7 Unknown (<1%) |
| **Race and Ethnicity** | |
| White | 4,137 (57%) |
| Black | 918 (13%) |
| Hispanic | 1,349 (18%) |
| Asian | 145 (2%) |
| Other | 759 (10%) |
| Not Reported | 8 (<1%) |
